# Chlamydiae in cnidarians: Shared functional potential despite broad taxonomic diversity

**DOI:** 10.1101/2023.11.19.567766

**Authors:** Justin Maire, Astrid Collingro, Matthias Horn, Madeleine J. H. van Oppen

## Abstract

Cnidarians, such as corals and sea anemones, associate with a wide range of bacteria that have essential functions, including nutrient cycling and the production of antimicrobial compounds. Within cnidarians, bacteria can colonize all microhabitats including the tissues. Among them are obligate intracellular bacteria of the phylum Chlamydiota (chlamydiae) whose impact on cnidarian hosts and holobionts remain unknown. Here, we conducted a meta-analysis of previously published cnidarian 16S rRNA gene metabarcoding data and eight metagenome-assembled genomes (MAGs) of cnidarian-associated chlamydiae to decipher their diversity and functional potential. While the metabarcoding dataset showed an enormous diversity of cnidarian-associated chlamydiae, five out of eight MAGs were affiliated with the Simkaniaceae family. The other three MAGs were assigned to the Parasimkaniaceae, Rhabdochlamydiaceae, and Anoxychlamydiaceae, respectively. All MAGs were associated with corals and showed a functional potential insufficient for an independent existence, lacking any nucleotide or vitamin and most amino acid biosynthesis pathways. Hallmark chlamydial genes, such as a type III secretion system, nucleotide transporters, and genes for host interaction, were encoded in all MAGs. Together these observations suggest an obligate intracellular lifestyle of cnidarian-associated chlamydiae. Cnidarian-associated chlamydiae lacked unique genes, suggesting the core chlamydial genetic arsenal may be flexible enough to infect many eukaryotic hosts, including cnidarians. Additional studies are needed to understand how chlamydiae interact with their cnidarian host, and other microbes in cnidarian holobionts. This first study of the diversity and functional potential of cnidarian-associated chlamydiae improves our understanding of both the cnidarian microbiome and the chlamydial lifestyle and host range.

## Introduction

The phylum Cnidaria comprises a wide range of early metazoans, including hard and soft corals, sea anemones, and jellyfish. These aquatic animals can have tremendous ecological value, such as scleractinian corals which form the foundation of coral reefs by building the reefs’ three-dimensional framework and driving its food web. Cnidarians associate with a wide range of microorganisms, which include protists, bacteria, archaea, viruses, and fungi [1–3]. Among them, bacteria have important functions, including protection against pathogens, and nitrogen and sulfur cycling [3–5]. Although the bacterial microbiome is highly diverse [6–8], most functional and genomic studies have focused on a few bacterial taxa, such as Proteobacteria, leaving most other taxa understudied. This includes the Chlamydiota (also known as chlamydiae), a phylum of obligate intracellular bacteria that infect a broad spectrum of eukaryotic hosts [9, 10]. All known chlamydiae have reduced genomes and a remarkably conserved biology; they are dependent on eukaryotic host cells, alternating between infectious extracellular elementary bodies and intracellular replicative reticulate bodies (Figure S1), and manipulate host cells through a type III secretion system (T3SS) [9]. While they are best known for being mammalian pathogens, chlamydiae, and especially members of the Simkaniaceae family, are regularly detected in metabarcoding data from scleractinian corals [11–14], octocorals [15, 16], cultures of intracellular dinoflagellates (Symbiodiniaceae) found in the tissues of corals, sea anemones, and jellyfish [17–19], as well as cultures of *Ostreobium* [20, 21], a green alga found in the skeleton of scleractinian corals.

Despite their prevalence, very little data exists on the diversity and function of cnidarian-associated chlamydiae, including whether they are pathogenic or mutualistic. This is likely due to their strictly host-associated lifestyle and generally low abundance in cnidarian holobionts. So far, only one study has analyzed the genome and location of a chlamydial associate of cnidarians: *Simkania* sp. Pac_F2b forms inclusions in the tentacles of the coral *Pocillopora acuta* and possesses all the chlamydial hallmark genes for host interaction, although its role remains unclear [12]. More data is therefore needed to increase our understanding of the taxonomic and genomic diversity of cnidarian-associated chlamydiae and their impact on cnidarian health. Here, we conducted a meta-analysis of previously published cnidarian microbiome data and genomes of cnidarian-associated chlamydiae. By analyzing their phylogeny and genomic content, we assessed whether chlamydiae detected in various cnidarian hosts and geographic locations represent distinct chlamydial lineages, share functional potential, and which convergent traits have evolved in these chlamydiae.

## Materials and methods

### Analysis of cnidarian microbiome data

Five databases were obtained from a previously published meta-analysis of 186 cnidarian and cultured Symbiodiniaceae microbiome studies [7]. The five datasets each group metabarcoding based on the hypervariable region of the 16S rRNA gene that was targeted (V1-V2, V3, V4, V5-V6, and V7-V8, respectively). The databases were analyzed in RStudio v2023.06.2 using the phyloseq package [22]. Amplicon sequence variants (ASVs) affiliating with the phylum Chlamydiota were identified in each dataset. Their relative abundance and prevalence (calculated as the percentage of samples within a family that harbors at least one chlamydial ASV) within cnidarian families and symbiodiniacean genera, as well as according to cnidarian health status was plotted in GraphPad Prism.

Chlamydial ASVs from each dataset were aligned to a reference alignment with SINA v1.7.2 [23] and subsequently clustered into genus-level OTUs with usearch v11.0.667 (-cluster_smallmem –id 0.95 -query_cov 0.9) [24]. The resulting datasets had the following OTUs: V1-V2 region: n = 4 (7 before clustering), V3: n = 363 (700 before clustering), V4: n = 939 (2638 before clustering), V5-V6: n = 367 (727 before clustering), V7-V8: n = 68 (96 before clustering). Centroid sequences of each cluster were then added to a chlamydial 16S rRNA gene reference multiple sequence alignment with MAFFT v7.5.20 (--addfragments – keeplength). The aligned centroid sequences for each dataset were then separately added to a chlamydial 16S rRNA gene phylogenetic tree with EPA-ng v0.3.8 [25] using the SYM+I+R10 model. The full-length 16S rRNA gene reference tree was based on a previously published phylogeny [26] and updated with full-length 16S rRNA gene sequences encoded in chlamydial MAGs published until 29.05.2023. The reference tree was inferred with IQ-TREE v2.2.5 [27] with 1000 ultrafast bootstrap replicates [28] and 1000 replicates of the SH-like approximate likelihood ratio test [29] under the SYM+I+R10 model. The resulting phylogenetic trees with added centroid sequences from the cnidarian datasets were rooted using Planctomycetota and Verrucomicrobiota sequences as outgroup and visualized with iTOL v6.8.1 [30].

### Phylogenetic analyses of metagenome-assembled genomes of cnidarian-associated chlamydiae

Eight metagenome-assembled genomes (MAGs) of cnidarian-associated chlamydiae were recovered from the GenBank/ENA/DDBJ database through the NCBI database query interface. The levels of completeness and contamination were assessed using CheckM2 v1.0.2 [31]. Taxonomic assignment was carried out using GTDB-Tk v2.1.0 [32] using the “classify_wf” workflow.

For comparative and phylogenetic analyses, a dataset of high quality chlamydial genomes based on Dharamshi et al. [33] was used. The chlamydial dataset was complemented by additional genomes available on GenBank/ENA/DDBJ on 29.05.2023. With few exceptions, only genomes with a completeness > 70%, contamination < 5% (both determined with CheckM2 v1.0.2 [31]), and average nucleotide identities < 95% (determined with FastANI v1.33 [34]) were considered for the final dataset, which included 169 chlamydial genomes and 89 genomes from Planctomycetota and Verrucomicrobiota serving as outgroup (Table S1).

To obtain the phylogenetic affiliation of the cnidarian-associated chlamydial MAGs, a set of 15 conserved non-supervised orthologous groups (NOGs) was used (Table S2). These 15 NOGs are known to retrieve the same topology for chlamydial phylogeny as the application of larger protein sets [33]. Proteins of the chlamydial and Planctomycetota-Verrucomicrobiota outgroup genomes belonging to the 15 NOGs were aligned with MAFFT v7.520 L-INS-i [34]. The resulting single protein alignments were subsequently trimmed with BMGE v2.0 [35] and concatenated. Four chlamydial MAGs with less than 50% of the NOGs present were excluded from phylogenetic analysis (Table S1). Maximum likelihood phylogeny was inferred with IQ-TREE v2.2.5 [27] with 1000 ultrafast bootstrap replicates [28] and 1000 replicates of the SH-like approximate likelihood ratio test [29] under the LG+F+I+R10 model. The resulting phylogenetic tree was rooted using the outgroup and visualized with iTOL v6.8.1 [30].

### Genome annotation and metabolic pathway reconstruction

Genome annotation was performed with Bakta v1.7.0 [36] and eggNOG-mapper v2.1.11 [37]. Metabolic pathways, transport systems and secretion systems were annotated via METABOLIC-G v4.0 [38] as previously described [39], applying “-m-cutoff 0.5” to include pathways which are ≥50% complete. The completeness of metabolic pathways (KEGG module database, https://www.genome.jp/kegg/module.html), transporters and secretion systems was estimated in EnrichM v0.6.4 [40]. KEGG modules of interest were then plotted in GraphPad Prism. The presence of specific genes of interest (virulence genes, T3SS) were investigated using BLAST and known sequences from the complete, high-quality genomes of either *Simkania negevensis* (unknown host) [41] or *Chlamydia trachomatis* (mammalian pathogen) [42] (Table S3). Secondary metabolites were predicted using antiSMASH v7.0.0 [43].

### Analysis of orthologous groups of proteins

For further comparative analysis of the chlamydial genomes (see Table S1), all encoded protein sequences were clustered into orthologous groups (OGs) with OrthoFinder v2.5.5 [44] under default parameters. OGs and their respective eggNOG annotations were merged in RStudio v2023.06.2 and analyzed. Protein sequences of six OGs were exclusively present in the genus *Simkania*, but did not show any sequence similarity to known proteins in public databases. Thus, the structures of representative sequences of these OGs were predicted with AlphaFold v2.3.2 [45] and visualized with UCSF ChimeraX v1.6.1 [46]. The resulting best-ranked protein structure models were then searched with structure search against RCSB protein data bank (PDB) [47]. Only hits with global pLDDT > 70 were considered in the results.

## Results

### Occurrence and prevalence of chlamydiae in cnidarians

By analyzing datasets from186 cnidarian and Symbiodiniaceae microbiome studies [7], we found that reads assigned to the phylum Chlamydiota are widespread in these host taxa, being detected in 1,925 individual samples out of 10,255, although usually at a low relative abundance (< 2% for all families except Psammocoridae, Figure 1). The highest relative abundance was in the scleractinian family, Psammocoridae (32% chlamydiae), although this data came from only one study (seven samples). The prevalence of chlamydiae was variable, though consistently high in certain scleractinian families, i.e., the Acroporidae (29% prevalence across 742 samples in the V4 region), Pocilloporidae (22% prevalence across 1128 samples in the V4 region), and Dendrophylliidae (72-75% prevalence across 136 samples in the V3, V4, and V5-V6 regions), and one actiniarian family, Aiptasiidae (24% prevalence across 501 samples in the V5-V6 region). In addition, chlamydiae were found in both healthy (0.14-2.39% across all cnidarians) and unhealthy (*i.e.*, stressed, bleached, damaged, or diseased; 0.01-1.97%) samples, often being slightly more abundant in healthy cnidarians (Figure S2). This shows that, while widespread, chlamydiae are not specifically associated with unhealthy or diseased cnidarians.

**Figure 1:**
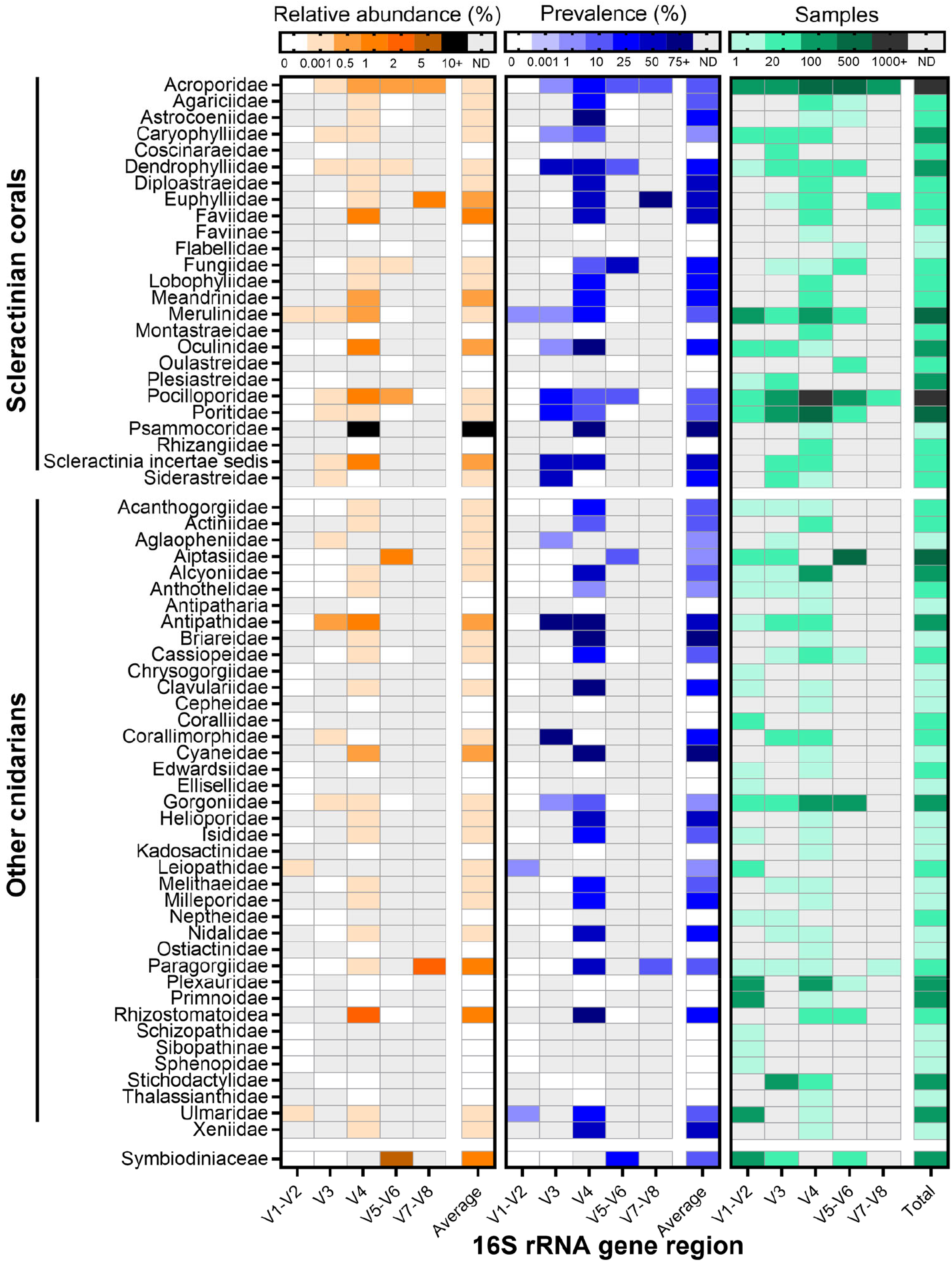
Relative abundance (left) and prevalence (middle) of chlamydiae amplicon sequence variants (ASVs) across cnidarian families and cultured Symbiodiniaceae. Prevalence was calculated as the number of samples per family that harbors at least one chlamydial ASV. The number of individual samples for each cell is also provided (right). Data was obtained from a meta-analysis of 186 16S rRNA gene metabarcoding studies [7]. ND: no data.

### Diversity of cnidarian- and Symbiodiniaceae**-**associated chlamydiae

To gain insights into the overall diversity of cnidarian- and Symbiodiniaceae-associated chlamydiae, we extracted the chlamydial ASVs from the metabarcoding datasets [7] and clustered the ASVs present in each dataset at genus-level identity (95%). The resulting cluster centroid sequences for each variable region of the 16S rRNA gene dataset were added to a chlamydial 16S rRNA gene reference tree. An enormous diversity of chlamydiae was detected in cnidarian and Symbiodiniaceae microbiomes (Figures 2 and S3-S6), with 1,741 genus-level chlamydial OTUs obtained from the five datasets. The vast majority of the ASVs affiliated with Simkaniaceae, Parasimkaniaceae and Rhabdochlamydiaceae. These families also contained the most abundant ASVs, though this may be impacted by the different sequencing depths of each study, as well as known biases from the most commonly used universal primers against chlamydiae (*e.g.* 515F/806R of the V4 region, or 1391R of the V8 region [48]). Only three chlamydial families were not found in cnidarians: Chlamydiaceae (only a handful of ASVs), Parilichlamydiaceae, Piscichlamydiaceae; these are all known to specifically infect chordate hosts.

**Figure 2:**
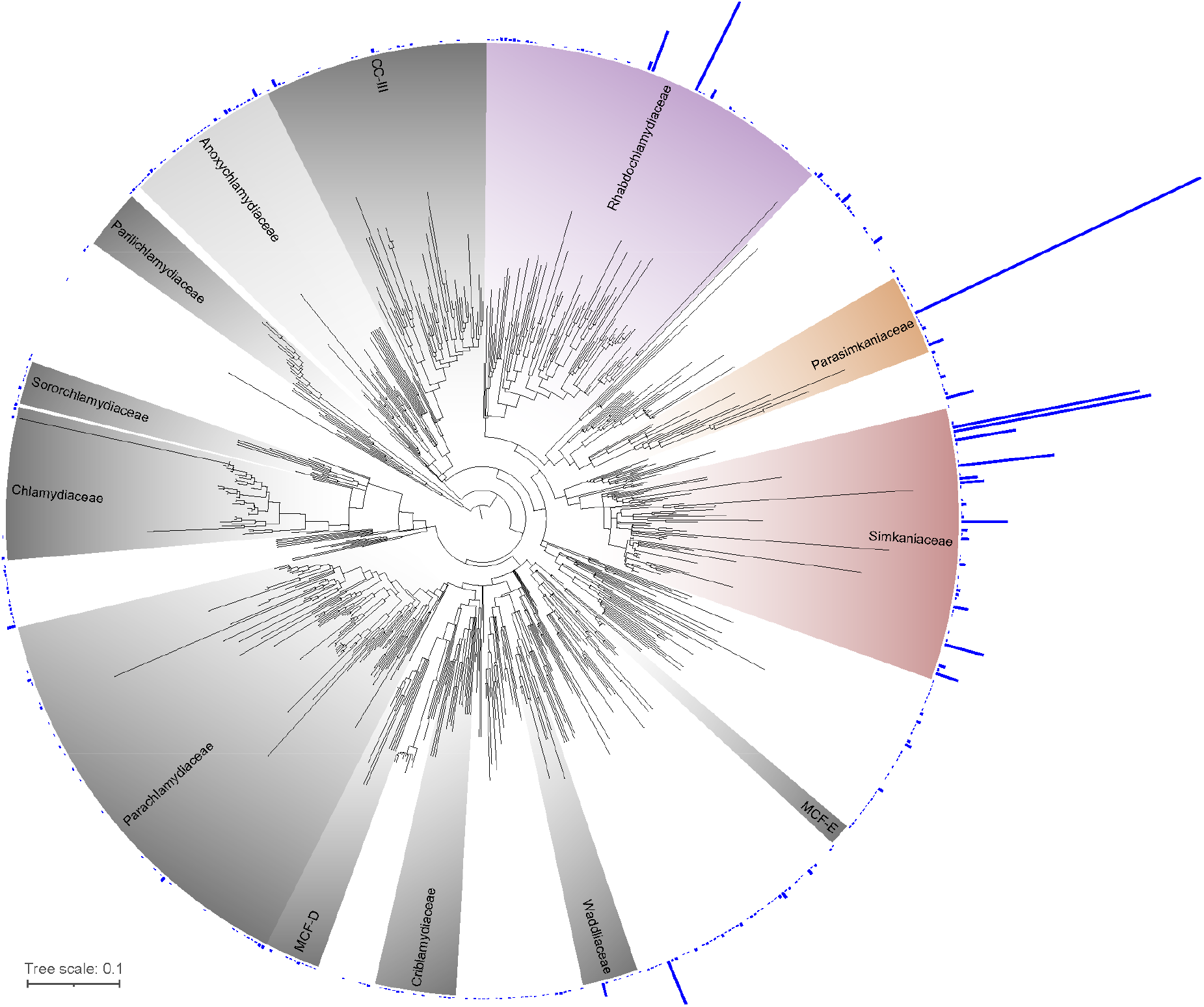
Diversity of cnidarian- and Symbiodiniaceae-associated chlamydiae. Maximum likelihood phylogeny of the 16S rRNA gene (V3 region) of 363 chlamydial genus-level OTUs. Blue bars represent the total read number (2-10,640 reads) found in a dataset of 43 cnidarian and Symbiodiniaceae 16S rRNA gene (V3 region) metabarcoding studies [7]. OTUs with no blue bars are reference sequences. Trees for four other 16S rRNA gene regions are available Figures S3-S6. Colored taxa (Simkaniaceae, Parasimkaniaceae, Rhabdochlamydiaceae, and Anoxychlamydiaceae) are families from which we obtained cnidarian-associated MAGs. MCF: metagenomic chlamydial family; CC-III: chlamydiae clade III.

### Metagenome-assembled genomes of cnidarian-associated chlamydiae

To better understand the nature of cnidarian-chlamydiae interactions, we analyzed eight metagenome-assembled genomes (MAGs) of cnidarian-associated chlamydiae detected previously in metagenomic studies (Tables 1 and S4). All MAGs were obtained from corals, one from an octocoral (Gorgoniidae, *Eunicella verrucosa*), and seven from scleractinian corals. MAG size ranged from 0.88 Mb to 1.88 Mb with estimated complete genome sizes ranging between 1.3 and 2 Mb. One MAG (EVH04_Bin2) was only 53.33% complete, while all others ranged from 78.84% to 100% in completeness. Contamination ranged from 0.07% to 3.14%. All but one (Pac_F2b [12]) were not analyzed in their original publication. To assess their taxonomy, we created a phylogenetic tree based on 15 conserved marker genes (Table S2), along with 161 other chlamydial genomes (Figure 3, Table S1). Five MAGs affiliated with members of the Simkaniaceae family, including four in the *Simkania* genus, according to GTDB-Tk classification, along with the type species *Simkania negevensis* and a MAG obtained from a Great Barrier Reef sponge (CLI4_bin_1 [49]). The other three MAGs belonged, respectively, to Rhabdochlamydiacaeae, Anoxychlamydiaceae, and the recently described Parasimkaniaceae, closely related to Simkaniaceae.

**Table 1:** Characteristics of eight published cnidarian-associated chlamydial MAGs. Additional details are available in Table S4.

| Genome ID | Host | Tissue | Location | Size (Mb) | Family | Study |
| --- | --- | --- | --- | --- | --- | --- |
| 3300010035-10 | *Cyphastrea* sp. | Unknown | Lord Howe Island, Australia | 1.32 | Simkaniaceae | [50] |
| CC_bin22 | *Acropora digitifera* | Surrounding seawater | Daya Bay, Shenzhen, China | 1.80 | Parasimkaniaceae | [51] |
| EVH04_Bin2 | *Eunicella verrucosa* | Unknown | Algarve, Pedra da Greta, Portugal | 0.88 | Rhabdochlamydiaceae | [52] |
| IP30a_bin.2 | *Isopora pallifera* | Skeleton | Heron Island, Australia | 1.06 | Simkaniaceae | [53] |
| Pac_F2b | *Pocillopora acuta* | Tentacle epidermis | Feather Reef, Australia | 1.25 | Simkaniaceae | [12] |
| PL23a_bin.60 | *Porites lutea* | Skeleton | Heron Island, Australia | 1.79 | Anoxychlamydiaceae | [53] |
| PL25a_bin.114 | *Porites lutea* | Skeleton | Heron Island, Australia | 1.88 | Simkaniaceae | [53] |
| PL25a_bin.130 | *Porites lutea* | Skeleton | Heron Island, Australia | 1.59 | Simkaniaceae | [53] |

**Figure 3:**
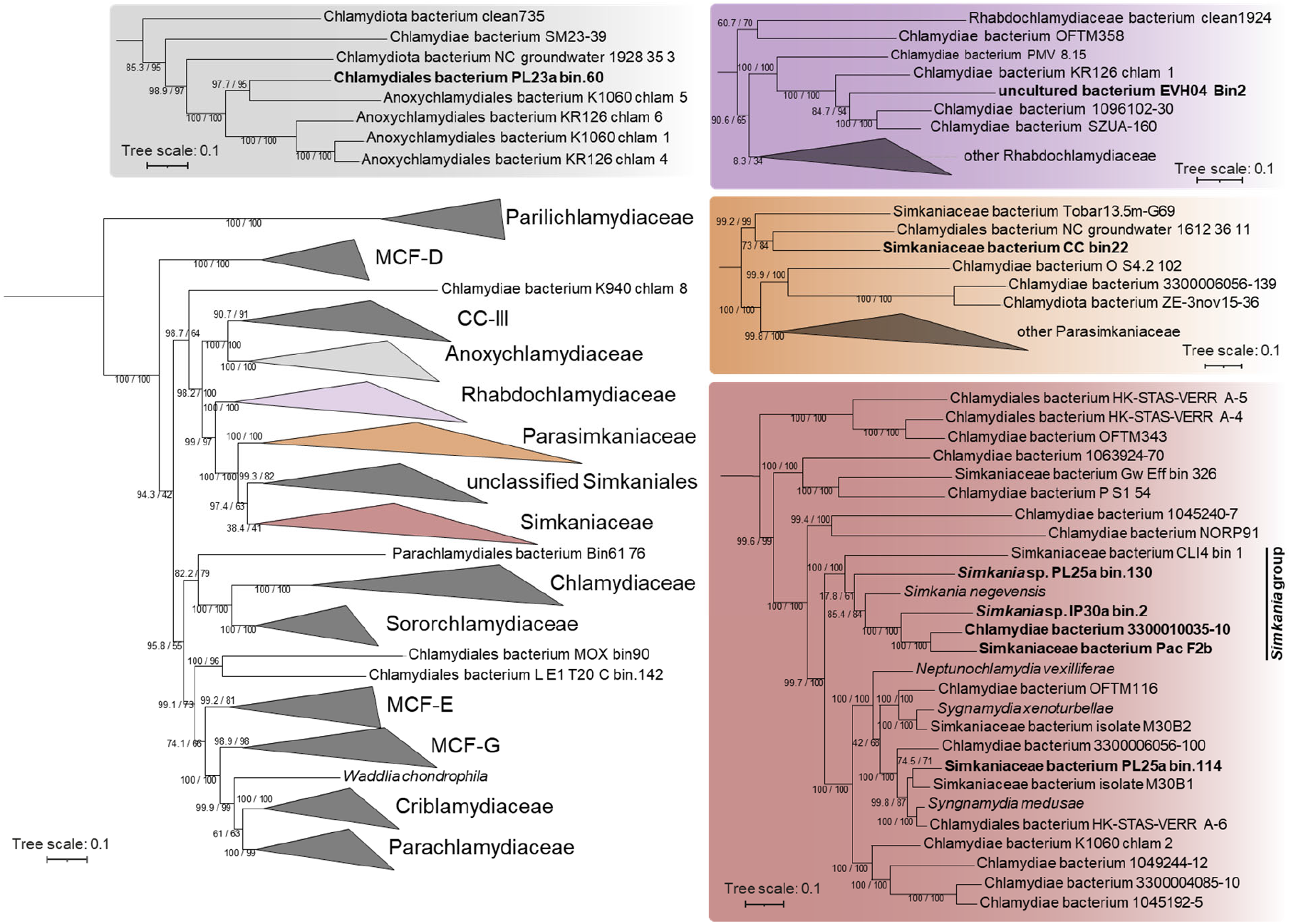
Chlamydial maximum likelihood phylogeny based on 15 conserved NOGs in 165 chlamydial genomes. Cnidarian-associated chlamydiae are in bold. Confidence values based on 1000 ultrafast bootstrap replicates and 1000 replicates of the SH-like approximate likelihood ratio test are provided. Additional data on the reference genomes is available in Table S1. MCF: metagenomic chlamydial family; CC-III: chlamydiae clade III.

### Functional potential of cnidarian-associated chlamydiae

Metabolic potential was assessed based on KEGG annotations and pathways (Figure 4). The complete genome of *S. negevensis* [41] was used as a reference. All MAGs showed reduced metabolic abilities, lacking pathways for the biosynthesis of nucleotides, vitamins, and most amino acids (only genes for glutamate, aspartate, lysine, and alanine biosynthesis were found, as well as glycine-serine interconversion, and the shikimate pathway). Most MAGs possessed the genes necessary for glycolysis, the tricarboxylic acid cycle, and the pentose phosphate pathway. The pathways were missing in EVH04_Bin2 (Rhabdochlamydiaceae) and IP30a_bin.2 (Simkaniaceae), which may be explained by their level of incompleteness (53.33% and 80.41% complete, respectively). The TCA was also missing from PL23a_bin.60 (Anoxychlamydiaceae), which instead possessed genes typically involved in anaerobiosis, such as genes encoding pyruvate:ferredoxin (flavodoxin) oxidoreductase, flavodoxin, rubredoxin, and the hydrogenase HydA.

**Figure 4:**
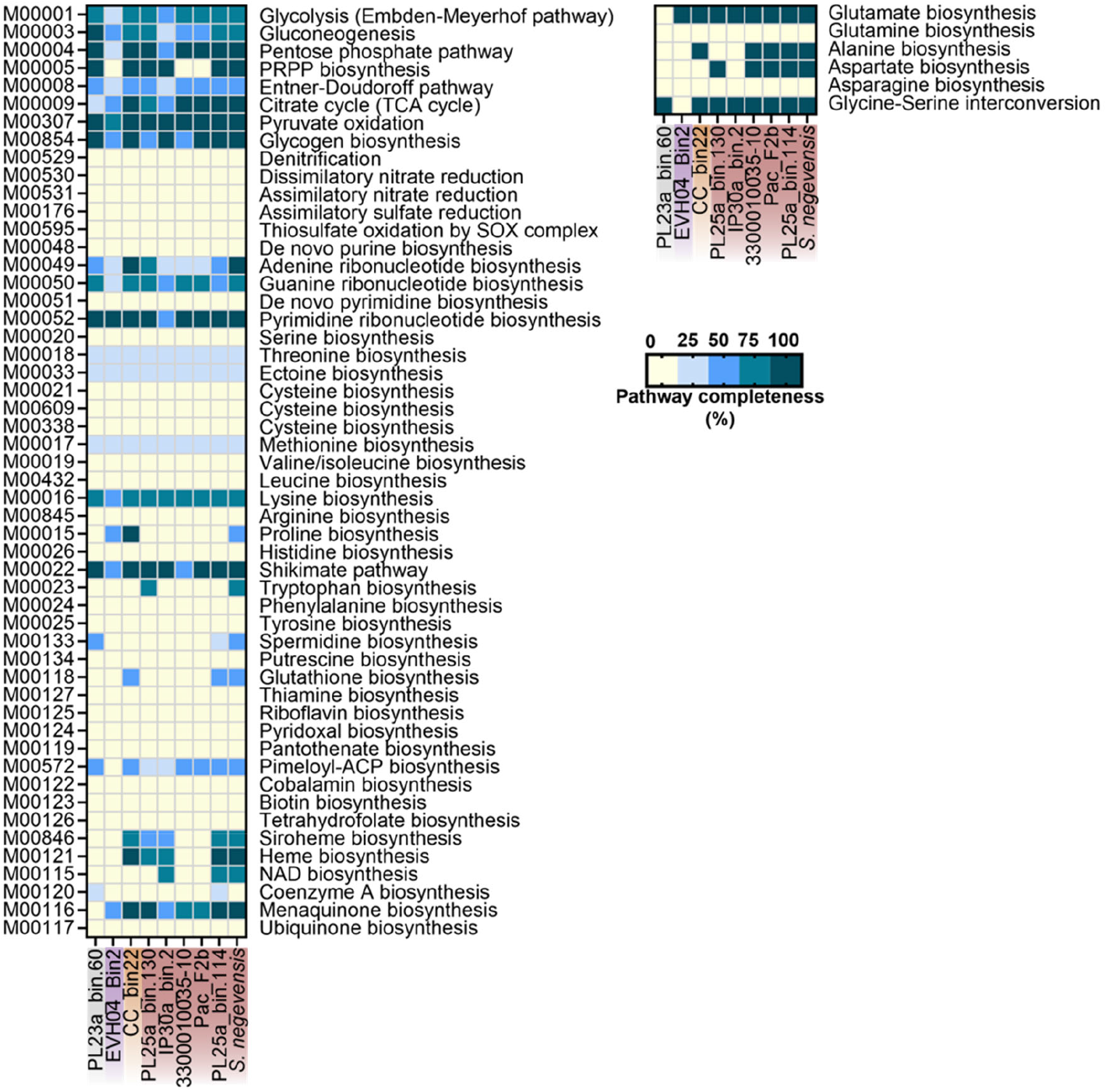
Metabolic potential encoded in eight cnidarian-associated chlamydial MAGs and *S. negevensis*. Genome names are colored according to the chlamydial family they belong to in Figure 3; grey: Anoxychlamydiaceae; purple: Rhabdochlamydiaceae; orange: Parasimkaniaceae; red: Simkaniaceae. The completeness of metabolic pathways (left panel; KEGG module database) of interest were annotated using METABOLIC-G and pathway completeness (in %) was estimated in EnrichM. The presence of genes involved in additional amino acid synthesis (right panel) was checked manually through the KEGG Reconstruct tool.

The pathways for the biosynthesis of two co-factors were complete in some MAGs: menaquinone (in five MAGs and *S. negevensis*, all part of the Simkaniaceae family) and heme (in four MAGs and *S. negevensis*, all part of the Simkaniaceae family). Yet, all ten genes of the heme biosynthesis pathway were missing from two other Simkaniaceae MAGs (3300010035-10 and Pac_F2b). Because of the large number of genes missing, this is unlikely a consequence of MAG incompleteness and it highlights a major difference between closely related Simkaniaceae. Finally, we wanted to infer the potential production of secondary metabolites and queried their presence in antiSMASH (Table S5). Four MAGs did not contain any predicted secondary metabolite gene cluster, while the other four (PL25a_bin.114, PL23a_bin.60, 3300010035-10, CC_bin22) possessed one each, all with putative antimicrobial properties. By contrast, the genome of *S. negevensis* was predicted to synthesize three secondary metabolites.

The pathogenic potential of cnidarian-associated chlamydiae was examined by similarity searches against hallmark chlamydial virulence-associated genes (Figure 5, Table S3) [54]. All MAGs had a complete T3SS, a key molecular machinery that allows for translocation of effectors into eukaryotic host cells, except for EVH04_Bin2 and IP30a_bin.2. However, none had genes to build a flagellar apparatus, which is at odds with other marine chlamydiae [55]. All MAGs encoded most chlamydial lifestyle- and virulence-associated genes that are present in other non-chordate infecting chlamydiae. This includes nucleotide transporters (ATP and nucleotide import), adhesins, T3SS effectors, Ser/Thr kinases (host cell modulation), and developmental regulators such as *euo*, which governs the transition between the two chlamydial developmental stages (Figure S1). Two genes involved in host cell lipid acquisition (*lpaT, aasC* [56]) were missing from one and three MAGs, respectively (EVH04_Bin2, IP30a_bin.2, and PL25a_bin.130), suggesting these MAGs may use a different mechanism for lipid acquisition. Finally, four potential T3SS effectors (*NUE,* CT_387, CT_783, and CT_830) were missing from 3-6 MAGs, with PL25a_bin.130 missing all four.

**Figure 5:**
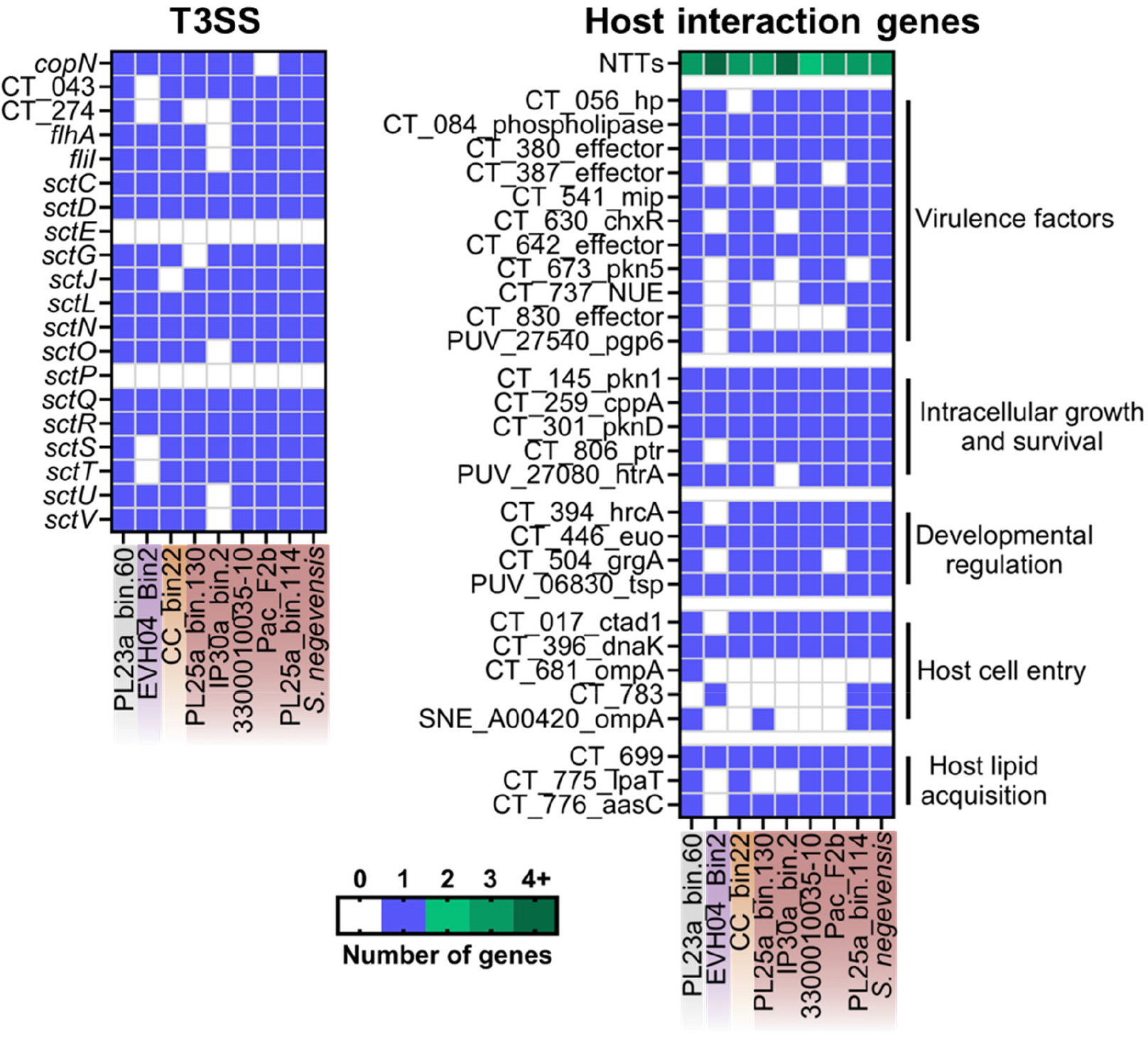
Important genes for host interactions encoded in cnidarian-associated chlamydial genomes and *S. negevensis*. Genome names are colored according to the chlamydial family they belong to in Figure 3; grey: Anoxychlamydiaceae; purple: Rhabdochlamydiaceae; orange: Parasimkaniaceae; red: Simkaniaceae. Genes important for host interactions from the complete genomes of *C. trachomatis, Parachlamydia acanthamoebae,* and *S. negevensis* were blasted against the eight coral-associated chlamydial MAGs to assess their presence. Additional information on the selected genes is available in Table S3.

### Genes unique to marine invertebrate-associated *Simkania*

To explore the presence of taxonomically restricted genes, we first queried the presence of genes specific to the eight coral-associated chlamydiae MAGs that are not found in other chlamydiae. No gene was found in all eight coral-associated chlamydial genomes that was not present in any of the other chlamydial genomes in the dataset (Table S1), which may be due to the taxonomic breadth represented by the eight coral-associated chlamydiae.

Therefore, we also analyzed the marine *Simkania* group, including one sponge-associated and four coral-associated *Simkania* (Figure 3). We found six genes that were present in at least four of the five MAGs and that were absent from all other chlamydiae genomes (Table 2). Because all six genes were predicted to encode hypothetical proteins without any known motifs, we predicted their structure using AlphaFold and queried the RCSB Protein Data Bank to identify structurally similar proteins (Table 2, Figure S7). Two out of six genes resulted in reliable structural predictions (pLDDT > 70) and had hits in the RCSB Protein Data Bank which also had structural predictions with a high confidence (pLDDT > 70). These included a peroxisomal biogenesis factor (involved in peroxisome formation) and an outer membrane protein.

**Table 2:**
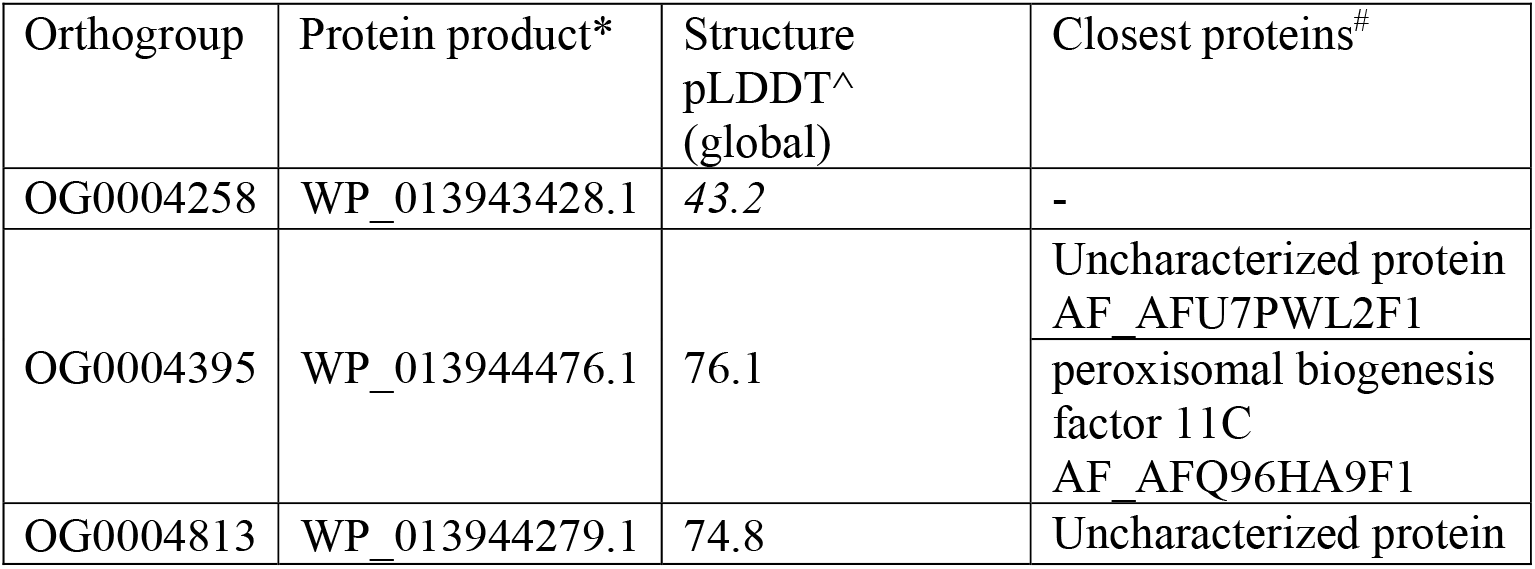
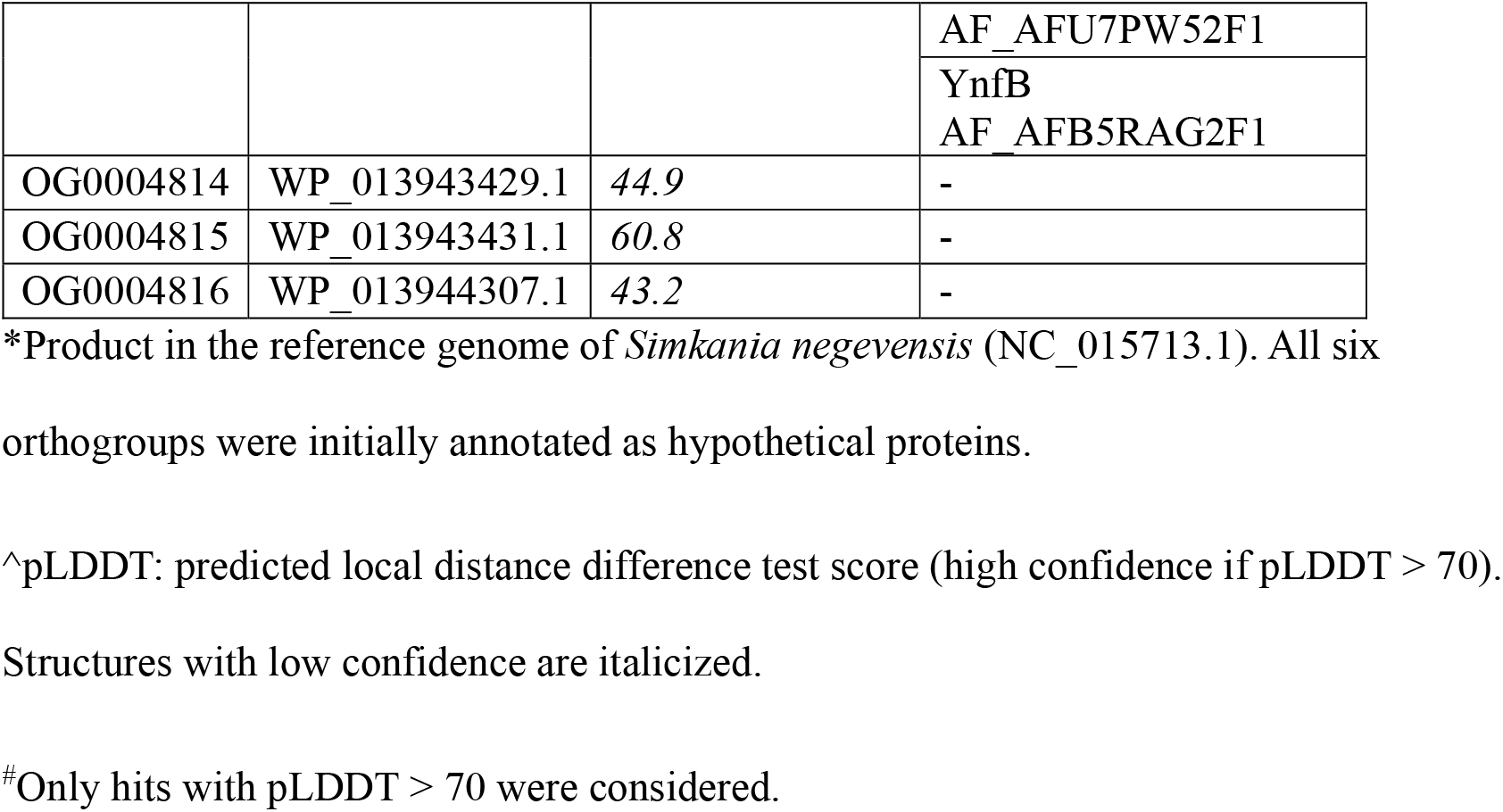
List of genes present in at least four out of the five MAGs of the marine *Simkania* group and absent from all other chlamydiae. The closest proteins based on structural predictions are provided.

## Discussion

Chlamydiae are among the most successful microbial symbionts of eukaryotes and infect a wide array of hosts, from protists to mammals [10]. This includes marine hosts and environments, with vast amounts of chlamydial sequences and genomes being retrieved from marine animals and sediments (chlamydiae may temporarily be extracellular as elementary bodies) [33, 54]. Despite being abundant and prevalent in cnidarians, cnidarian-associated chlamydiae have remained largely understudied. Here, we analyzed previously published data from cnidarian and Symbiodiniaceae microbiome studies and eight chlamydial MAGs sequenced from cnidarian hosts (corals), to increase our understanding of both chlamydial and cnidarian biology, and lay the foundations for further studies on cnidarian-chlamydiae interactions. The sequences derived from 186 cnidarian and Symbiodiniaceae microbiome studies [7] enabled a comprehensive overview of the presence, prevalence, and diversity of chlamydiae in cnidarian holobionts. Cnidarian-associated chlamydiae were highly diverse; with the exception of members of the chordate-infecting Piscichlamydiaceae, Parilichlamydiaceae, and Chlamydiaceae, members of all other chlamydial families occur in cnidarian microbiomes. High chlamydial diversity has also been described for sponges [33], pointing towards potentially deep evolutionary relationships with these two groups of early metazoans.

Six out of eight MAGs were members of the Simkaniaceae and Parasimkaniaceae families, which are often found to be highly abundant in cnidarian metabarcoding studies [7, 11, 14]. The Simkaniaceae family contains many marine representatives derived from jellyfish, sponges, fish, and marine sediments [33, 54], with a recent meta-analysis showing that around 60% of published Simkaniaceae 16S rRNA sequences originated from marine environments, with another ∼30% from animal hosts (host environment unspecified) and engineered environments (*e.g.*, bio-reactors, wastewater) [54]. This shows that marine ecosystems harbor many suitable hosts for Simkaniaceae. Strikingly, four cnidarian-associated MAGs, along with a sponge-associated MAG, are close relatives of the type species *S. negevensis* and likely belong to the *Simkania* genus. *S. negevensis* was initially discovered as a laboratory contaminant of human cell cultures in Israel [57], although its original host was never elucidated. Its phylogenetic relatedness with coral- and sponge-associated Simkaniaceae highlighted in our study points to a marine origin of *S. negevensis*.

Two additional cnidarian-associated MAGs belonged to the Rhabdochlamydiaceae and Anoxychlamydiaceae. The Anoxychlamydiaceae MAG (PL23a_bin.60) was obtained from coral skeleton, a few centimeters underneath the algal (*Ostreobium)* layer [53]. The skeleton is anoxic past the first few centimeters [58], and probably up to the coral tissue layer at night, when phototrophic endoliths (*e.g., Ostreobium*, Cyanobacteria) do not photosynthesize [59], and could therefore support anaerobic microorganisms such as Anoxychlamydiaceae [60]. This is consistent with the incompleteness of the TCA cycle, an aerobic process, in PL23a_bin.60 and the presence of genes encoding pyruvate:ferredoxin (flavodoxin) oxidoreductase, flavodoxin and rubredoxin observed also in other Anoxychlamydiaceae and are consistent with anaerobic metabolism [60].

Comparative gene content analysis did not detect any genes that are restricted to coral-associated chlamydiae. A similar conclusion was drawn from a recent analysis of sponge-associated chlamydiae, which found that gene content was related to phylogenetic affiliation rather than to an ecological association with their host [33]. Broader metabolic pathway reconstruction corroborated this lack of specificity, as most coral-associated chlamydiae shared metabolic and host interaction abilities with chlamydiae from other hosts. Within the marine *Simkania* group, however, the comparative analysis identified six genes that are not found in any other chlamydiae. Based on structural predictions with AlphaFold, functions could be hypothesized for two of the six genes. OG0004395 may represent peroxisomal biogenesis factor-like proteins, which could interfere with host peroxisome size and fission. Eukaryotic peroxisomes play a role in clearing bacterial infections by assisting with phagocytosis [61]. For instance, *C. trachomatis* interacts with host peroxisomes, possibly for phospholipid synthesis [62]. Marine *Simkania* may therefore interact with host peroxisomes for cell growth or phagocytosis escape. OG0004813 includes proteins most closely related to bacterial outer membrane proteins, and thus these proteins may be involved in host cell adhesion. A key difference among the marine *Simkania* was that the heme biosynthesis pathway was completely absent from two out of four *Simkania* MAGs. This stark difference in closely related organisms may be explained by factors unrelated to chlamydial or host taxonomy, such as the host environment (*e.g.,* depth, light conditions, temperature variability) and which host microhabitat the *Simkania* reside in (*e.g.,* gastrodermis, epidermis, skeleton). Mutations in the *hemG* gene, involved in heme synthesis, were associated with increased infectivity in *C. trachomatis* [63], suggesting that the presence or absence of the heme biosynthesis pathway may lead to different infection outcomes. Despite this difference, the overall gene content similarities among marine *Simkania* are consistent with previous studies finding that chlamydiae in general have a large core genome [41, 54, 64], with similar metabolic abilities and lifestyle features, such as energy parasitism and a biphasic developmental cycle. The lack of recognizable host-driven genomic convergence in several chlamydial groups may provide more host flexibility and explain the wide range of eukaryotic hosts they have successfully formed associations with.

While genomic analyses confirmed that cnidarian-associated chlamydiae possess all the genes needed for the chlamydial intracellular, parasitic lifestyle, it remains unclear whether they are beneficial or detrimental to cnidarians, as it is in many other organisms, and if their presence is an indicator of holobiont health. We found that chlamydiae prevalence was similar between healthy and unhealthy cnidarians, and no cnidarian disease is known to be caused by chlamydiae. Bleached cnidarians did not show increased chlamydial abundance either, suggesting chlamydiae may not opportunistically infect stressed animals, as seen for example with the coral pathogen *Vibrio coralliilyticus* [65]. Thus, as for sponges [33], negative impacts of chlamydiae infections in cnidarians have not been observed. Potentially beneficial functions, however, remain elusive, with the exception of the production of secondary metabolites which may be involved in chemical defense against other microorganisms. They could therefore function as defensive symbionts, as recently shown in protist hosts where chlamydiae provide protection against *Legionella* [66, 67] or giant viruses [68]. This would however require every cell of exposed tissue to harbor chlamydiae. Functional studies should be conducted, where holobiont fitness metrics must be compared between infected and uninfected colonies of the same cnidarian species and geographic location (*e.g., Acropora loripes* from Davies Reef, Australia [11]). In combination with gene expression analyses of both host and chlamydiae, through metatranscriptomics or spatial transcriptomics, such experiments may shed light on cnidarian-chlamydiae interactions. Finally, while chlamydiae are abundantly found in cnidarian metabarcoding studies, cnidarian samples often contain symbiotic microeukaryotes, such as fungi and protists (*e.g.*, Symbiodiniaceae, *Ostreobium*, corallicolids, chromerids) [2, 69]. Chlamydiae have been detected in laboratory cultures of both Symbiodiniaceae and *Ostreobium* [17–21]. Whether chlamydiae can infect these protists *in hospite* remains unknown. Four MAGs analyzed in our study were isolated from the coral skeleton [53], and may therefore be derived from *Ostreobium* or other endolithic protists. One MAG (Pac_F2b) was obtained from bacterial clusters excised from coral epidermis and the strain represented by this MAG therefore likely infects coral cells. Protist-associated chlamydiae (*e.g.* members of the Amoebachlamydiales) have larger genomes (∼2-3Mb) compared to animal-associated chlamydiae (∼1-2Mb) [41, 64]. Larger genomes may offer more flexibility to infect unicellular organisms, which provide less stable conditions than multicellular organisms. The MAGs analyzed in our study were all smaller than 2 Mb, suggesting they are coral-rather than protist-associated. Further studies combining genomic and spatial analyses are needed to fully elucidate the hosts and functions of chlamydiae within cnidarian holobionts [8, 12, 13, 70, 71].

In conclusion, we provide here the first genomic meta-analysis of cnidarian-associated chlamydiae. We uncovered that chlamydiae are highly prevalent in a wide range of cnidarians across vast spatial scales. Cnidarian-associated chlamydiae are highly similar to other chlamydiae and we found little genomic specialization. Additional functional studies and broader sequencing of cnidarian microbiomes will be needed to fully understand the breadth of cnidarian-chlamydiae interactions, including where they sit on the mutualism-parasitism continuum and whether they are an indicator of cnidarian health. Our research adds to the immense diversity of chlamydial hosts already described and improves our understanding of chlamydial evolution, and the broader evolution of intracellular symbioses.

## Acknowledgments

We thank the Melbourne Research Cloud (University of Melbourne) and the Life Science Compute Cluster (University of Vienna) for providing the high-performance computing instances needed for this work.

## Data availability statement

All data used in this study was previously published (See Tables S1 and S4 for accession numbers).

## Author contribution

Conceptualization: JM, AC, MH, MvO; Formal analysis: JM, AC; Visualization: JM, AC; Funding acquisition: JM, AC, MH, MvO; Writing - Original Draft Preparation: JM; All authors reviewed and edited the final manuscript.

## Notes

### Competing Interest Statement

The authors have declared no competing interest.

