## Supplementary material for "Chlamydiae in cnidarians: Shared functional potential despite broad taxonomic diversity"

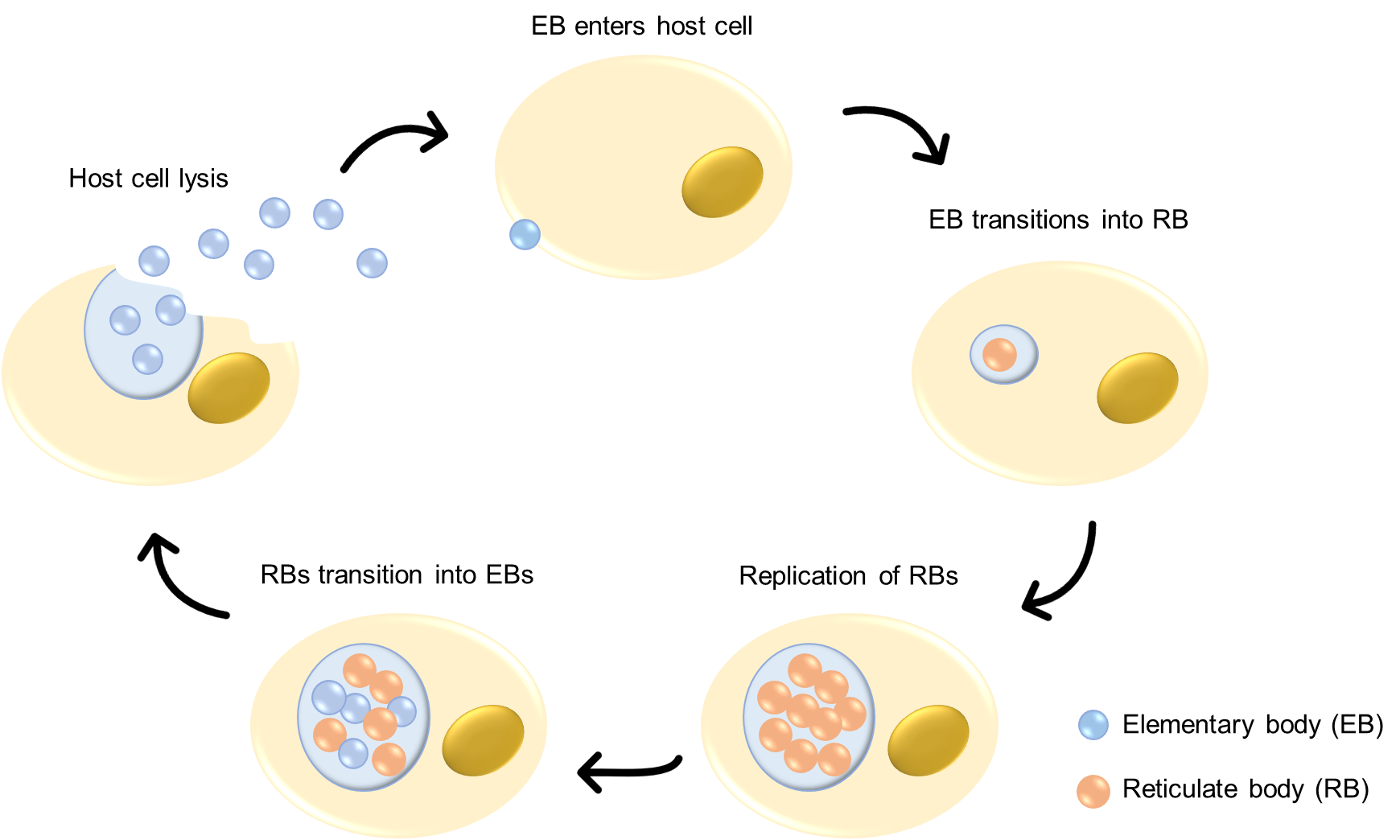

**Figure S1:** The chlamydial lifecycle. Chlamydiae alternate between infectious extracellular elementary bodies (EBs) and intracellular replicative reticulate bodies (RBs).

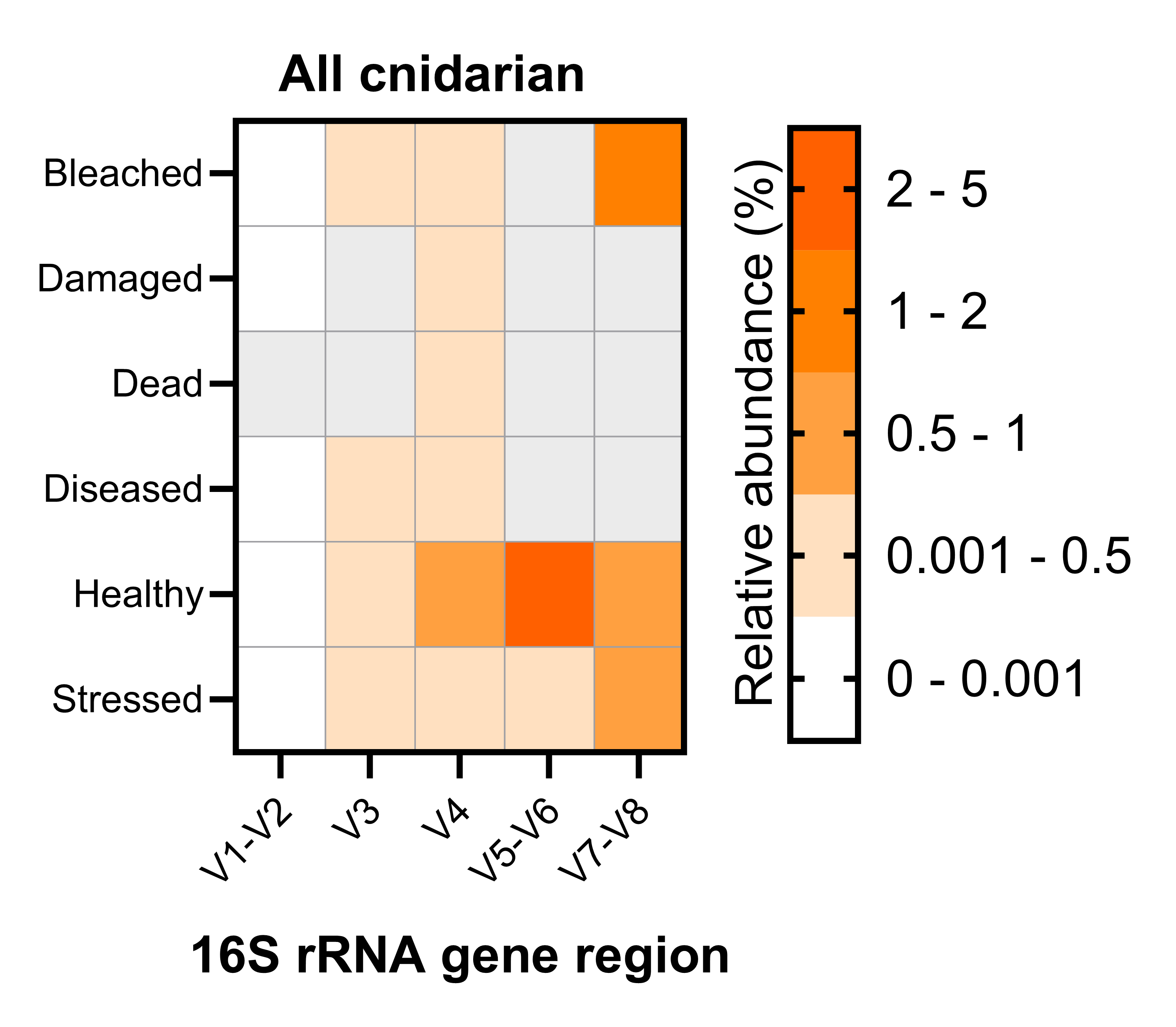

**Figure S2:** Relative abundance of chlamydiae amplicon sequence variants (ASVs) across all cnidarian families, according to host health status. Data was obtained from published databases regrouping almost 200 cnidarian 16S rRNA gene metabarcoding studies [7].

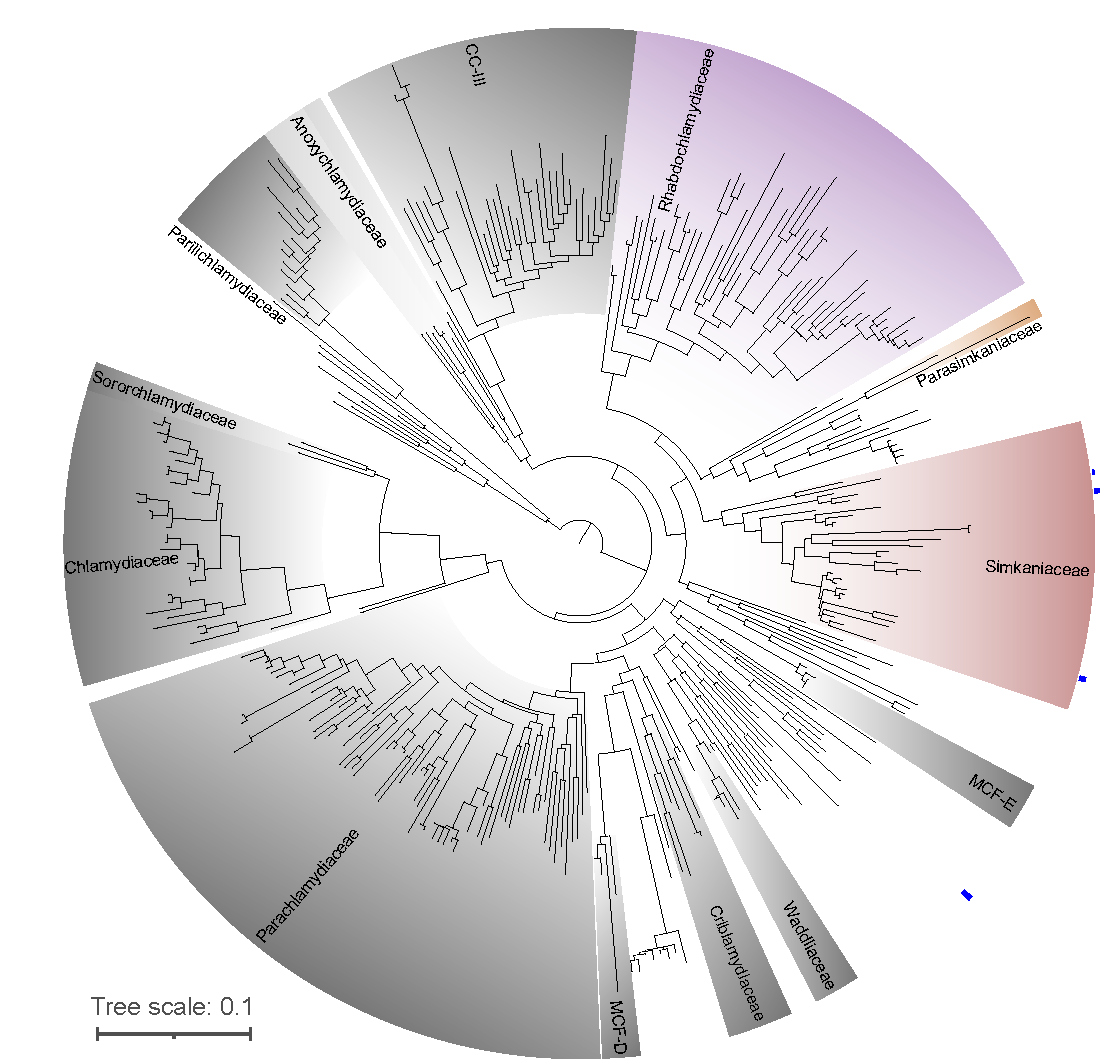

**Figure S3:** Maximum likelihood phylogeny of the 16S rRNA gene (V1-V2 region) of four cnidarian- and Symbiodiniaceae-associated chlamydial genus-level OTUs. Blue bars represent the total read number (2-47 reads) found in a dataset of 19 cnidarian and Symbiodiniaceae 16S rRNA gene (V1-V2 region) metabarcoding studies [6]. OTUs with no blue bars are reference sequences. Colored taxa (Simkaniaceae, Parasimkaniaceae, Rhabdochlamydiaceae, and Anoxychlamydiaceae) are families from which we obtained cnidarian-associated MAGs. MCF: metagenomic chlamydial family; CC-III: chlamydiae clade III.

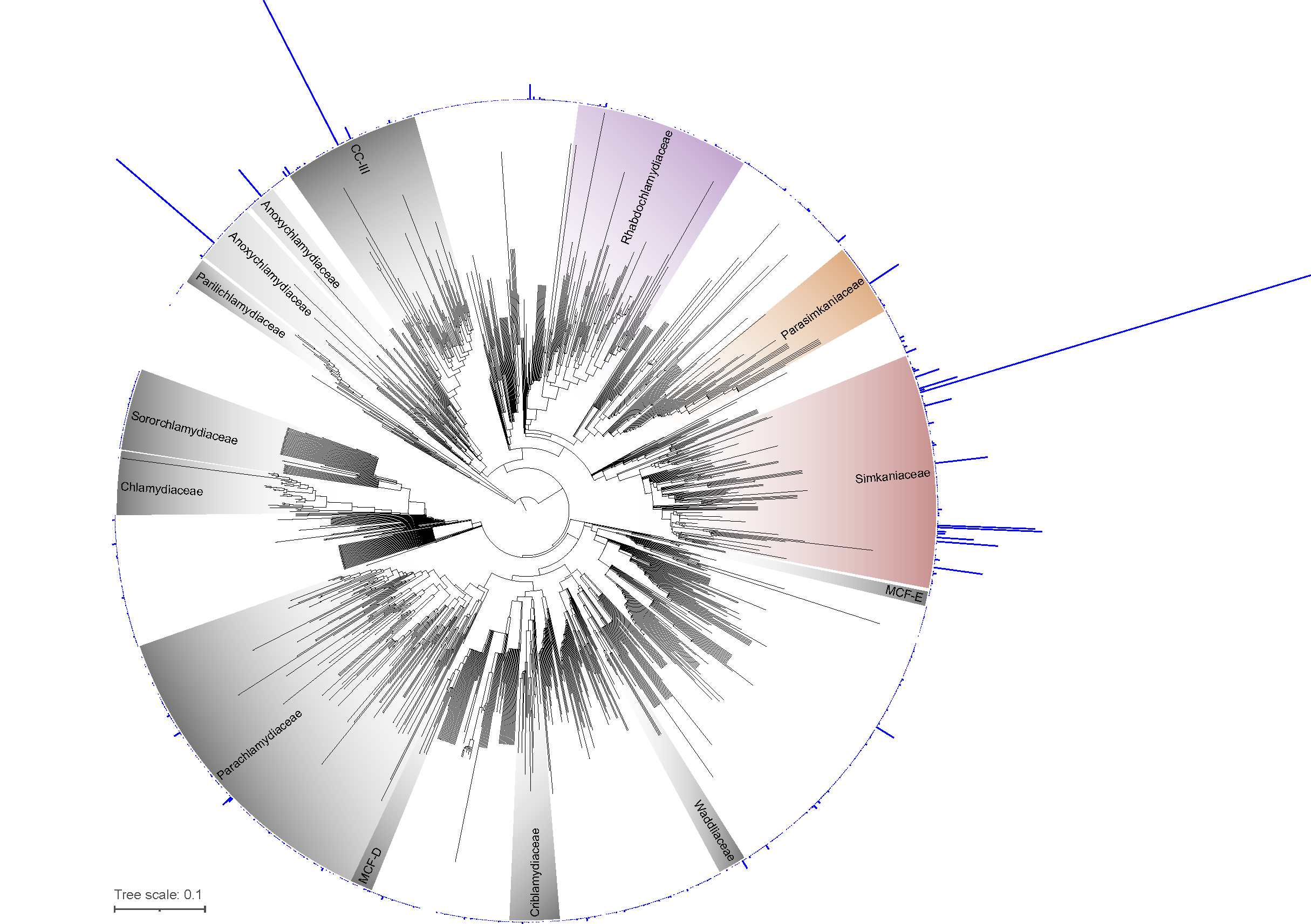

**Figure S4:** Maximum likelihood phylogeny of the 16S rRNA gene (V4 region) of 939 cnidarian- and Symbiodiniaceae associated chlamydial genus-level OTUs. Blue bars represent the total read number (2-132,585 reads) found in dataset of 85 cnidarian and Symbiodiniaceae 16S rRNA gene (V4 region) metabarcoding studies [6]. OTUs with no blue bars are reference sequences. Colored taxa (Simkaniaceae, Parasimkaniaceae, Rhabdochlamydiaceae, and Anoxychlamydiaceae) are families from which we obtained cnidarian-associated MAGs. MCF: metagenomic chlamydial family; CC-III: chlamydiae clade III.

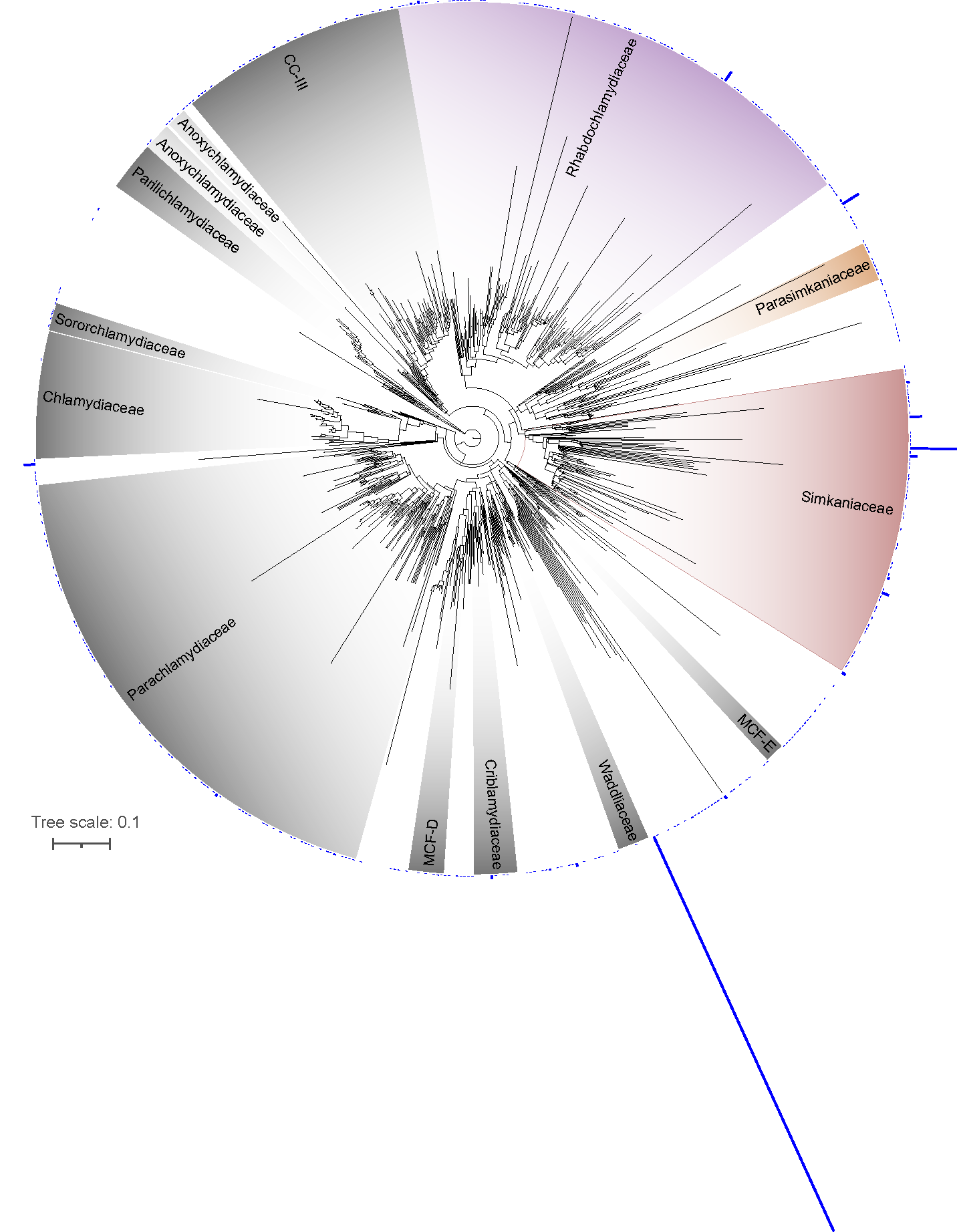

**Figure S5:** Maximum likelihood phylogeny of the 16S rRNA gene (V5-V6 region) of 367 cnidarian- and Symbiodiniaceae-associated chlamydial genus-level OTUs. Blue bars represent the total read number (2-339,497 reads) found in a dataset of 44 cnidarian and Symbiodiniaceae 16S rRNA gene (V5-V6 region) metabarcoding studies [6]. OTUs with no blue bars are reference sequences. Colored taxa (Simkaniaceae, Parasimkaniaceae, Rhabdochlamydiaceae, and Anoxychlamydiaceae) are families from which we obtained cnidarian-associated MAGs. MCF: metagenomic chlamydial family; CC-III: chlamydiae clade III.

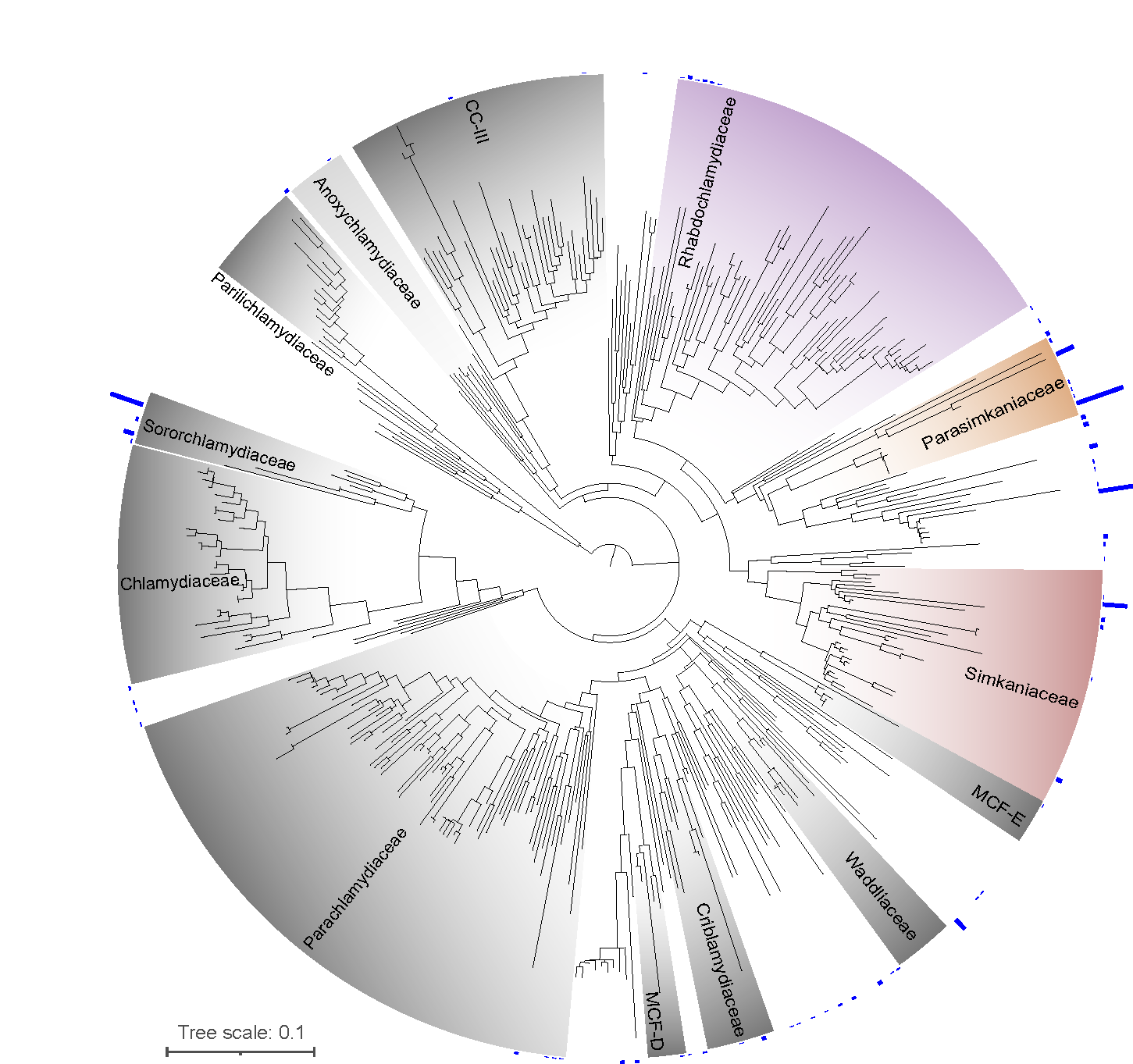

**Figure S6:** Maximum likelihood phylogeny of the 16S rRNA gene (V7-V8 region) of 68 cnidarian- and Symbiodiniaceae-associated chlamydial genus-level OTUs. Blue bars represent the total read number (2-2,128 reads) found in a dataset of 9 cnidarian and Symbiodiniaceae 16S rRNA gene (V7-V8 region) metabarcoding studies [6]. OTUs with no blue bars are reference sequences. Colored taxa (Simkaniaceae, Parasimkaniaceae, Rhabdochlamydiaceae, and Anoxychlamydiaceae) are families from which we obtained cnidarian-associated MAGs. MCF: metagenomic chlamydial family; CC-III: chlamydiae clade III.

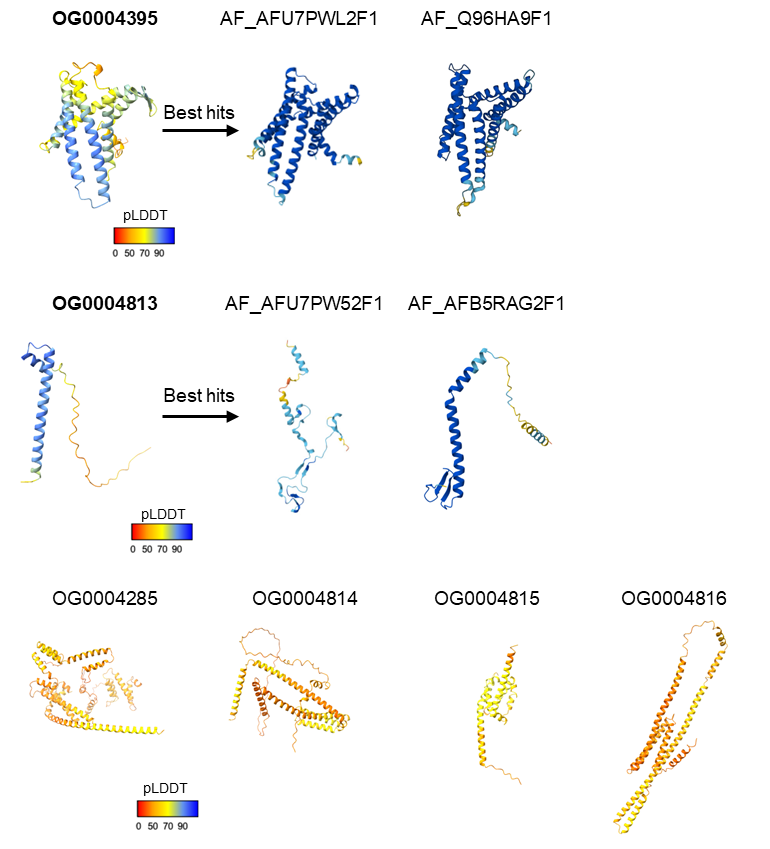

**Figure S7:** Predicted structures of six orthogroups unique to the marine *Simkania* group (see Table 2 for more details). Only structures for OG0004395 and OG0004813 (in bold) were of high enough confidence (global pLDDT > 70). The two closest hits for each of them are also provided.

**Table S1:** List of chlamydial genomes used for phylogenetic analyses.

(attached)

**Table S2:** Marker Non-supervised Orthologous Group (NOG) proteins used for chlamydial phylogenetic analysis.

| NOG | NOG category | NOG description |
| --- | --- | --- |
| COG0064 | J | Aspartyl-tRNA (Asn)/glutamyl-tRNA (Gln) amidotransferase subunit B |
| COG0092 | J | Ribosomal protein S3 |
| COG0233 | J | Ribosome recycling factor |
| COG0290 | J | Translation initiation factor IF-3 |
| COG0292 | J | Ribosomal protein L20 |
| COG0323 | L | DNA mismatch repair protein MutL |
| COG0335 | J | Ribosomal protein L19 |
| COG0342 | U | Preprotein translocase subunit SecD |
| COG0468 | L | DNA recombination/repair protein RecA |
| COG0532 | J | Translation initiation factor IF-2 |
| COG0536 | DL | GTPase Obg involved in cell cycle, chromosome segregation and ribosome assembly |
| COG0706 | M | Membrane protein insertase YidC |
| COG1185 | J | Polyribonucleotide nucleotidyltransferase Pnp |
| COG1530 | J | Ribonuclease G or E |
| COG1663 | M | Tetraacyldisaccharide-1-P 4’-kinase LpxK |

**Table S3:** List of the chlamydiae genes queried in this study (Figure 4), related to type III secretion system (T3SS) components (A) or other lifestyle and virulence processes (B), and their putative functions in host-chlamydiae interactions.

(attached)

**Table S4:** Detailed characteristics of eight published MAGs belonging to cnidarian-associated chlamydiae.

(attached)

**Table S5:** List of predicted secondary metabolites in cnidarian-associated chlamydiae and *Simkania negevensis*.

(attached)
